# A Key Role for *S-*Nitrosylation in Immune Regulation and Development in the Liverwort *Marchantia polymorpha*

**DOI:** 10.1101/2025.09.29.679193

**Authors:** Nadra Tabassum, Justin Goodrich, Gary J. Loake

## Abstract

Nitric oxide (NO) is an important signaling molecule in flowering plant immunity. It rapidly accumulates in response to pathogen perception. In addition to it’s direct response to microbes, NO controls a range of defence responses primarily through *S*-nitrosylation. This process is a redox-dependent modification where a NO group attaches to the thiol of a cysteine residue, creating an *S*-nitrosothiol (SNO). To explore the role of *S*-nitrosylation more broadly, we characterised the single-copy *S*-*nitrosoglutathione reductase 1* (Mp*GSNOR1*) gene in the liverwort *Marchantia polymorpha* (Marchantia), a representative of a lineage widely diverged from flowering plants. We generated loss-of-function alleles using CRISPR/Cas9 genome editing. Disrupting Mp*GSNOR1* resulted in pronounced morphological alterations, highlighting the role of GSNOR1 in the structural development of Marchantia. Additionally, we show that Mp*GSNOR1* is essential for SNO homeostasis and immune function. Our results suggest that GSNOR was part of the tool kit of the ancestral land plant and functioned in immunity and development.

**Highlight:** First evidence from a Liverwort shows GSNOR controls immunity and development via *S*-nitrosylation, revealing these regulatory roles as ancient traits of land plants.

## Introduction

The transition of plants from water to land about 470 to 515 million years ago significantly influenced the evolution of global biodiversity (Bowman et al., 2016, 2022; Romani et al., 2024). This transition is thought to have driven the evolution of a highly efficient and specialised plant immune system to protect against potential infection from co-evolving microbial pathogens (Carella et al., 2019; Poveda, 2020). Successful plant migration from water to land probably relied on the capacity of these plants to deal with various microorganisms that had already been successfully adapted to terrestrial conditions (Matsui et al., 2019; Yotsui et al., 2023).

Evolutionary molecular plant-microbe interactions (EvoMPMI) is a rapidly emerging field that has contributed to advancing our knowledge regarding the emergence of early plant immune systems (Gimenez-Ibanez et al., 2019; Redkar et al., 2022). In this regard, *Marchantia polymorpha* (hereafter Marchantia) is showing promise as an excellent model species for comparative studies to characterise evolution of traits and ancestral functions (Bowman et al., 2017, 2022; Alcaraz et al., 2018).

Marchantia is named after the French botanist Nicolas Marchant and has been studied for its life cycle and medicinal properties since the 16th century (Bowman et al., 2016). Interest in Marchantia as a model species has been rekindled both by technical advances such as the availability of a genome sequence and facile gene disruption via genome editing as well as by its phylogenetic position as an extant representative of a lineage that diverged from that of flowering plants at least 450 million years ago. It is readily transformed and has a relatively small genome of 250 Mb with low redundancy for many of its developmental and regulatory genes (Bowman et al., 2017; Iwasaki et al., 2021; Montgomery et al., 2020). Marchantia features a dominant haploid gametophyte generation in its life cycle, facilitating gene discovery and characterisation through forward and reverse genetic screens (Shimamura, 2016). It reproduces both asexually, via clonal propagules termed gemmae, and sexually via different male (antheridophore) and female (archegoniophore) structures born on separate male and female individuals (Bowman, Araki and Kohchi, 2016; Shimamura, 2016). It has nine chromosomes, including a sex specific U or V chromosome that determines female or male identity, respectively (Shimamura, 2016).

Recent research has demonstrated that several phytopathogens, including *Xylaria cubensis, Colletotrichum species, Phytophthora palmivora, Pseudomonas syringae* (*Pst*), *Fusarium oxysporum* (Fo) and several fungal endophytes can grow well within and thrive in Marchantia. Visibly apparent symptoms of infections by these pathogens are generally composed of disease symptoms like necrosis and chlorosis on the thallus surface (Carella et al., 2018; Redkar et al., 2022)Click or tap here to enter text. Recently, the study on the interaction of Marchantia and various *Trichoderma* species has revealed a range of results, including enhanced resistance, growth promotion, or increased disease susceptibility (Poveda et al., 2023). The study also studied patterns of tissue colonisation and found that tissue colonisation additionally relates to transcription activation of certain genes related to the defence system, forming such products as salicylic acid (SA)(Poveda et al., 2023). The expression of these functional responses depends on a conserved immune repertoire. This includes LysM-domain pattern-recognition receptors (MpLYK1 and MpLYR), nucleotide binding leucine-rich repeat (NLR)-like resistance factors, and parts of the SA pathway, which suggests that important elements of the land plant immune system likely existed in early diverging lineages. (Matsui et al., 2019; Yotsui et al., 2023).

However, the immune structure of this liverwort is still mostly unexplored (Bowman et al., 2022). Recent research has highlighted the role of the thallus of Marchantia as a mediator of susceptibility (Carella et al., 2018; Iwakawa et al., 2021). In particular, the air chambers, which help with gas exchange, may serve as entry points for pathogens. Marchantia Mp*nop1* mutants, which do not develop these chambers, show greater resistance to *P. palmivora, Pst*, and *Agrobacterium tumefaciens*, indicating that a lack of access to internal tissues does not allow their colonisation (Carella et al., 2018; Iwakawa et al., 2021; Tsuboyama-Tanaka & Kodama, 2015).

Interrogation of the Marchantia genome has uncovered twenty-three resistance (R)-like NLR genes (Bowman et al., 2017), which in angiosperms encode proteins that predominantly recognise the activity of pathogen effector proteins (Bowman et al., 2017; Lin et al., 2012). In *P. palmivora* infections, in planta expression of virulence genes, including RxLR effectors, proteases, and cell wall-degrading enzymes, has been observed, along with the formation of haustoria, highlighting pathogen strategies for immune evasion and host manipulation (Carella et al., 2018). In the case of *Pst*, mutants deficient in type III secretion (*hrcC–)* display significantly reduced virulence, while effectors like avrPto and avrPtoB suppress host immunity, mirroring their roles in flowering plants (Gimenez-Ibanez et al., 2019).

In addition, Marchantia also produces the key immune signalling molecule SA in response to attempted pathogen infection. Further, the genome of Marchantia possesses homologs of all critical components associated with SA biosynthesis and signalling found in vascular plants(Gimenez-Ibanez et al., 2019; Lin et al., 2012). By contrast, algal genomes only include select components of the SA molecular machinery, indicating that this system may have evolved during plant land colonisation (Bowman et al., 2017; Gimenez-Ibanez et al., 2019).

In vascular plants, a key feature of the immune response following recognition of potential pathogens is the engagement of a rapid nitrosative burst, leading to the accumulation of nitric oxide (NO) and numerous other reactive nitrogen intermediates (RNIs) (Yu et al., 2014). These small redox-active molecules regulate a plethora of key immune-related functions, including the oxidative burst (Yun et al., 2011), SA-based signalling (Loake & Grant, 2007; Moreau et al., 2010; Tada et al., 2008), and immune-related transcriptional reprogramming (Cui et al., 2018; Imran et al., 2018). This regulation, through the transfer of NO bioactivity predominantly occurs by *S*-nitrosylation, the reversible attachment of an NO moiety to a rare, highly reactive protein cysteine (Cys) thiol (SH), to form an *S*-nitrosothiol (SNO) (Stomberski et al., 2019; Xia et al., 2014; Yu et al., 2014). Thus, this redox-based, post-translational modification is akin to more well-established post-translational modifications (PTMs) such as phosphorylation (Cohen, 2002; Sefton & Shenolikar, 1995), in the control of protein function.

The extent of cellular *S*-nitrosylation can be controlled directly by Thioredoxin h5 (Trxh5) through the SNO reductase activity of this enzyme (Kneeshaw & Spoel, 2018) and indirectly, via the turnover of *S*-nitrosoglutathione (GSNO), which functions as a storage reservoir for NO bioactivity (Kneeshaw & Spoel, 2018; Yun et al., 2016), by GSNO reductase (GSNOR) (Chen et al., 2009; Feechan et al., 2005; Lee et al., 2008). This enzyme is required for the resistance gene mediated protection and systemic acquired resistance (Feechan et al., 2005; Tada et al., 2008; Wang, 2006; Wang et al., 2013). Furthermore, the absence of *GSNOR1* inhibited the accumulation of SA and resulted in delayed and reduced expression of SA-dependent genes in Arabidopsis (Feechan et al., 2005). *GSNOR1* function is therefore required for both SA biosynthesis and associated signal transmission (Feechan et al., 2005; Tada et al., 2008; Wang, 2006; Wang et al., 2013).

Thus, while the pathogen-triggered nitrosative burst and cognate redox signalling is a key feature of the immune response in flowering plants, a potential role for this molecular machinery in the immune systems of other land plant lineages remains to be established. To address this deficiency, we identified an orthologue for *GSNOR* in Marchantia and subsequently generated a series of loss-of-function alleles for this gene by CRISPR/Cas9-based gene editing.

Similar to the phenotypes observed in vascular plant *GSNOR* mutants (Kwon et al., 2012; Lee et al., 2008), disruption of Mp*GSNOR* affected not only pathogen defense but also fundamental developmental processes. Mutants displayed reduced thallus growth, defective gemma cup and rhizoid formation, delayed sporophyte development, and increased susceptibility to *Pst* DC3000 infection. These findings suggest that redox homeostasis, particularly through GSNOR-mediated NO signaling, involved in redox homeostasis during development and immunity, is conserved between the widely diverged lineages of liverworts and flowering plants. This implies the mechanism was likely present in the most recent common ancestor of land plants as part of its ancestral signaling network.

## Materials and Methods

### Plant material and growth condition

The male accession Takaragaike-1 (Tak-1) of *Marchantia* was used as the wild-type (WT) in this study. Gemmae of Marchantia were cultivated in vitro on half-strength Gamborg’s B5 medium supplemented with 1% agar (solid medium) or without agar (liquid medium). Cultures were maintained under a 16-h light/8-h dark photoperiod, with a light intensity of 110 µmol m⁻² s⁻¹, at 20°C and a relative humidity of 60–65%. For bacterial infection experiments, 2-to 4-week-old plants were transferred to controlled environmental chambers maintained at 22°C (range, 16–24°C), 45–65% relative humidity, and short-day conditions (8 hours of light).

### Phylogenetic analysis

The Arabidopsis GSNOR1 protein sequence was used in BLASTP searches to retrieve similar proteins from various databases as follows: Marpolbase (https://marchantia.info) for Marchantia, *Physcomitrella patens, Selaginella moellendorffii, Azolla filiculoides, Amborella trichopoda, Spirogloea muscicola, Chlamydomonas reinhardtii.* Genbank (https://blast.ncbi.nlm.nih.gov/Blast.cgi) for *Cyanidioschyzon merolae and Picea sitchensis.* The *Cyanophora paradoxica* sequence was retrieved from the transcriptome ast the Cyanophora genome project (http://cyanophora.rutgers.edu/cyanophora_v2018/).

Sequences with E values less that 10^-42^ were selected and aligned using MAFFT implemented within the Geneious package (https://www.geneious.com) using AUTO algorithm and BLOSUM62 scoring matrix. The alignment was manually trimmed, and positions with >30% gaps were stripped using the mask alignment tool in Geneious. The model selection tool within IQTree (http://iqtree.cibiv.univie.ac.at/) was first used to select the LG + R5 model and the maximum likelihood tree inferred with IQ-TREE. The output tree was exported as a .svg file and edited using FigTree (http://tree.bio.ed.ac.uk/software/figtree/) and Inkscape (https://inkscape.org/).

### Generation of CRISPR/Cas9-mediated Mp*gsnor1* knockout mutants

Mp*gsnor1* knockout mutants in Marchantia were generated using CRISPR/Cas9 technology through *Agrobacterium*-mediated gemma (Agar Trap) transformation of Tak-1(WT) gemmae (Tsuboyama et al., 2018; Tsuboyama & Kodama, 2014). Two guide RNAs (MpGNR_sgRNA1 and MpGNR_sgRNA2), Supplementary File Table S1, targeting the second exon of Mp*GSNOR1* were designed using CRISPRdirect (https://crispr.dbcls.jp) and cloned into the pGE_En03 entry vector as per Sugano et al. (2018). The constructs were recombined into the Cas9-containing binary vector pMpGE011 and introduced into *Agrobacterium tumefaciens* GV3101 (mp90). Two independent mutant alleles, Mp*gsnor1-1^ge^* and Mp*gsnor1-^4ge^*, were recovered. We maintained the primary transformants (G₁ generation) as stable lines, and gemmae produced by these G₁ plants (G₂ generation) were collected and used for all subsequent phenotypic and molecular examinations.

Transformed lines were screened by PCR using gene-specific primers that amplified ∼459 bp fragments spanning the target site (primer sequences are listed in Table S1). Agarose gel electrophoresis was used to detect larger insertions or deletions. However, since many CRISPR-induced mutations were small indels undetectable by gel, a subset of transformants was randomly selected for Sanger sequencing to confirm the presence of mutations, particularly frameshifts likely to disrupt gene function. Sequence alignments against the wild-type Mp*GSNOR1* sequence were used to verify edits. Although the Cas9 transgene was not eliminated, periodic sequencing was used to verify the stability of the mutations across generations.

### Generation of Complementation constructs

For complementation experiments in Marchantia, constructs expressing Arabidopsis *GSNOR1* (At*GSNOR1*) were generated using Gateway cloning technology. The At*GSNOR1* coding sequence was amplified from Arabidopsis cDNA using primers listed in Supplementary file, Table-S1, cloned into the pDONR™221 vector (Invitrogen) by BP clonase mediated recombination according to the manufacturer’s instructions, and verified by sequencing. Following recombination by Gateway LR Clonase II enzyme (Life Technologies), the At*GSNOR1* sequences were inserted downstream of a *35S* promoter in the Gateway-compatible binary vector pMpGWB102. Constructs were introduced into *Agrobacterium tumifaciens* strain GV3101 (mp90) by heat shock transformation and subsequently used to transform the Mp*gsnor1-1^ge^*and Mp*gsnor1-4^ge^* mutant backgrounds by G-Agar trap (Gemma transformation) methods (Tsuboyama et al., 2018). The presence of the T-DNA construct in the putative transformants were confirmed by PCR using At*GSNOR1-*specific primers (Table S1). The complementation line *pro35S:*At*GSNOR1^ge^* in Mp*gsnor1-4^ge^* background was subsequently used for the experiments.

### Transformation of *Marchantia*

Transformation of Marchantia was performed using the G-agar trap method (Tsuboyama & Kodama, 2018). Transformants were selected on half-strength Gamborg’s B5 medium containing 1% agar, 100 µg/ml cefotaxime, and 0.5 µM chlorsulfuron to isolate non-chimeric transgenic gemmae. For complementation of the Mp*gsnor1* mutants, selection was performed on half-strength Gamborg’s B5 medium containing 1% agar, 100 µg/ml cefotaxime, and 10 µg/ml hygromycin to recover super transformants (carrying both the Cas9 gene editing T-DNA and the At*GSNOR1* construct T DNA).

### Bacterial strains and inoculation methods

*Pseudomonas syringae* pv. *tomato* DC3000 (*Pst* DC3000) was cultivated on Luria-Bertani (LB) medium supplemented with rifampicin (50 µg/ml) at 28°C. Overnight cultures were grown at 28°C with shaking at 300 rpm. Inoculation methods followed the procedure described by Gimenez-Ibanez et al. 2019. Briefly, Marchantia gemmalings were grown on Whatman filter paper over half-strength Gamborg’s B5 agar medium at 21°C under a 16-h light/8-h dark cycle. Plants were inoculated by dipping in a bacterial suspension of 10□ CFU/ml (OD600=0.2) containing 0.04% Silwet L-77 for 5 minutes. Post-inoculation, plants were transferred to soil under short-day conditions. Bacterial growth was quantified by plating serial dilutions of thallus disc homogenates harvested 3-or 5-days post-inoculation.

### GSNOR activity assay

GSNOR activity was determined by monitoring NADH consumption at 340 nm, following established methods (Feechan et al., 2005). Protein extracts (100 µg) from wild-type and mutant plants were incubated at 25°C in a 1 ml reaction mixture containing 20 mM Tris-HCl (pH 8.0), 0.2 mM NADH, and 0.5 mM EDTA. Reactions were initiated by adding *S*-nitrosoglutathione (GSNO) at a final concentration of 300 µM. Activity was expressed as nmol NADH consumed per minute per mg of protein.

### Biotin-switch assay for total *S*-nitrosylation measurement

The assay was performed essentially as described by Jaffrey & Snyder (2001) with modifications for Marchantia. Four-week-old Marchantia plants were inoculated with *Pst* DC3000 or mock-treated with 10 mM MgCl₂. Approximately 200 mg of thallus tissue (fresh or stored at –80 °C) was ground to a fine powder in liquid nitrogen and homogenised with four volumes of extraction buffer (250 mM HEPES, 1 mM EDTA, 0.1 mM neocuproine, pH 7.7, supplemented with 0.5% (v/v) Triton X-100 and 1× protease inhibitor cocktail). Homogenates were divided into 1.5–2 ml microcentrifuge tubes and centrifuged at 14,000 rpm (∼20,000 × g) for 15 min at 4 °C. The supernatant was filtered through a 0.45 µm syringe filter, and protein concentration was determined by absorbance at 280 nm (NanoDrop). Samples were normalised to 200 µg protein in HEN buffer (250 mM HEPES, 1 mM EDTA, 0.1 mM neocuproine, pH 7.7) and aliquoted into 100 µl volumes. For free thiol blocking, 300 µl blocking buffer (HEN containing 5% SDS and 50 mM N-ethylmaleimide (NEM) was added to each sample. Tubes were wrapped in aluminum foil and incubated at 50 °C for 20 min with intermittent vortexing. Proteins were precipitated by adding two volumes of ice-cold 100% acetone and incubating at –20 °C for at least 20 min (or overnight). Pellets were collected by centrifugation (14,000 rpm, 5 min, 4 °C), washed three times with 70% (v/v) acetone at 4 °C, and air-dried briefly in the dark. Pellets were resuspended in 96 µl HENS buffer (HEN containing 1% SDS), and freshly prepared sodium ascorbate (500 mM, 12 µl) and biotin-HPDP (12 µl, final 0.4 mM; Thermo Fisher) were added. Reactions were incubated at room temperature for 1–2 h on a rocker, protected from light. Control reactions were done without sodium ascorbate. After labelling, 120 µl 2× SDS loading buffer was added, and samples were loaded onto SDS–PAGE gels without boiling. Following electrophoresis and transfer to PVDF membranes, S-nitrosylated proteins were detected using HRP-conjugated anti-biotin antibody (#7075, Cell Signaling Technology) and visualised by chemiluminescence.

### RNA extraction and cDNA synthesis

Total RNA was extracted from approximately 100 mg of three-week-old Marchantia thalli using TRIzol (Invitrogen) followed by purification with Qiagen spin columns according to the manufacturer’s protocol. RNA quality and concentration were assessed by agarose gel electrophoresis and NanoDrop spectrophotometry, respectively. cDNA synthesis was performed using either the High-Capacity cDNA Reverse Transcription Kit (Thermo Fisher Scientific) for isolating the Arabidopsis GSNOR1 cDNA or the LunaScript RT SuperMix Kit (NEB #E3010) for qRT-PCR analyses following the guidelines of manufacturer.

### Quantitative real-time PCR (qRT-PCR)

qRT-PCR reactions were performed using Luna Universal qPCR Master Mix (NEB #M3003). Each 20 µl reaction included 10 µl master mix, 0.5 µl forward and reverse primers (10 µM each), and 1 µl cDNA template. Reactions were carried out in a Roche LightCycler 480 instrument under the following cycling conditions: initial denaturation at 95°C for 60 s, followed by 45 cycles at 95°C (15 s) and 60°C (30 s). Melt curve analysis (60–95°C) verified amplification specificity. Relative gene expression was calculated using the ΔΔCt method with Mp*ACT* (Marchantia *Actin*) as a reference gene (Saint-Marcoux et al., 2015). Three biological and three technical replicates were performed per treatment. Primer sequences are provided in Supplementary file, Table S1.

### Differential interference contrast (DIC) microscopy

Morphological features of thalli and rhizoids were visualized using a Nikon Eclipse E600 microscope equipped with differential interference contrast (DIC) illumination and polarisation filters. Samples were mounted on glass microscope slides and examined at magnifications of 10×, 20×, or 40×. Images were captured using an attached digital camera, and scale bars were added using the accompanying camera software for accurate size referencing.

### Statistical Analysis

All statistical analyses were conducted utilizing GraphPad Prism (version 10.0; GraphPad Software, Inc., San Diego, CA). Phenotypic alterations, GSNOR activity, biotin switch assay outcomes, qRT-PCR expression data, and pathogenicity assessments were analysed employing unpaired two-tailed Student’s t-tests, as indicated in the figure legends and the corresponding number of replicates for each experiment. Results were considered statistically significant at P < 0.05.

## RESULTS

### Marchantia has a single *GSNOR1* orthologue

GSNOR1 is a member of the class III alcohol dehydrogenases (ADHs) distinguished by its use of GSNO as substrate and NADH as cofactor. BLASTP searches using Arabidopsis At*GSNOR1* (At5g43940) as query against the Marchantia proteome retrieved the Mp1g16170 product (hereafter MpGSNOR1) as the best match. The two proteins showed very high similarity throughout their lengths (81% identity over 374 amino acids). Several other Marchantia proteins also had high similarity over the full length of At*GSNOR1* – for example, the Mp2g02600 and Mp8g16300 products had 53 and 51% identity, respectively. To explore relationships further, we made a maximum likelihood phylogenetic analysis (Figure 1) sampling GSNOR1 and other class III ADHs from diverse plant and algal lineages. This resolved a very strongly supported clade that contained GSNOR1 orthologues from all the major lineages of the Archaeplastida (plants, green algae, red algae, and glaucophytes) and confirmed that Mp*GSNOR1* is the only Marchantia orthologue – in other words, MpGSNOR1 is more closely related to GSNOR1 from Arabidopsis or algae, etc, than it is to Mp2g02600 or any other Marchantia protein. The other Marchantia proteins with similarity to GSNOR1 mapped elsewhere in the phylogeny, suggesting that they are ADH class enzymes but unlikely to use GSNO as substrate. Consistent with this, multiple sequence alignments (Figure S1) revealed that although the residues involved in coordinating the structural and catalytic zinc atoms are conserved in all the Marchantia proteins, only in Mp*GSNOR1* were the residues involved in contacting the NAD(H) cofactor and GSNO substrate well conserved.

**Figure 1.**
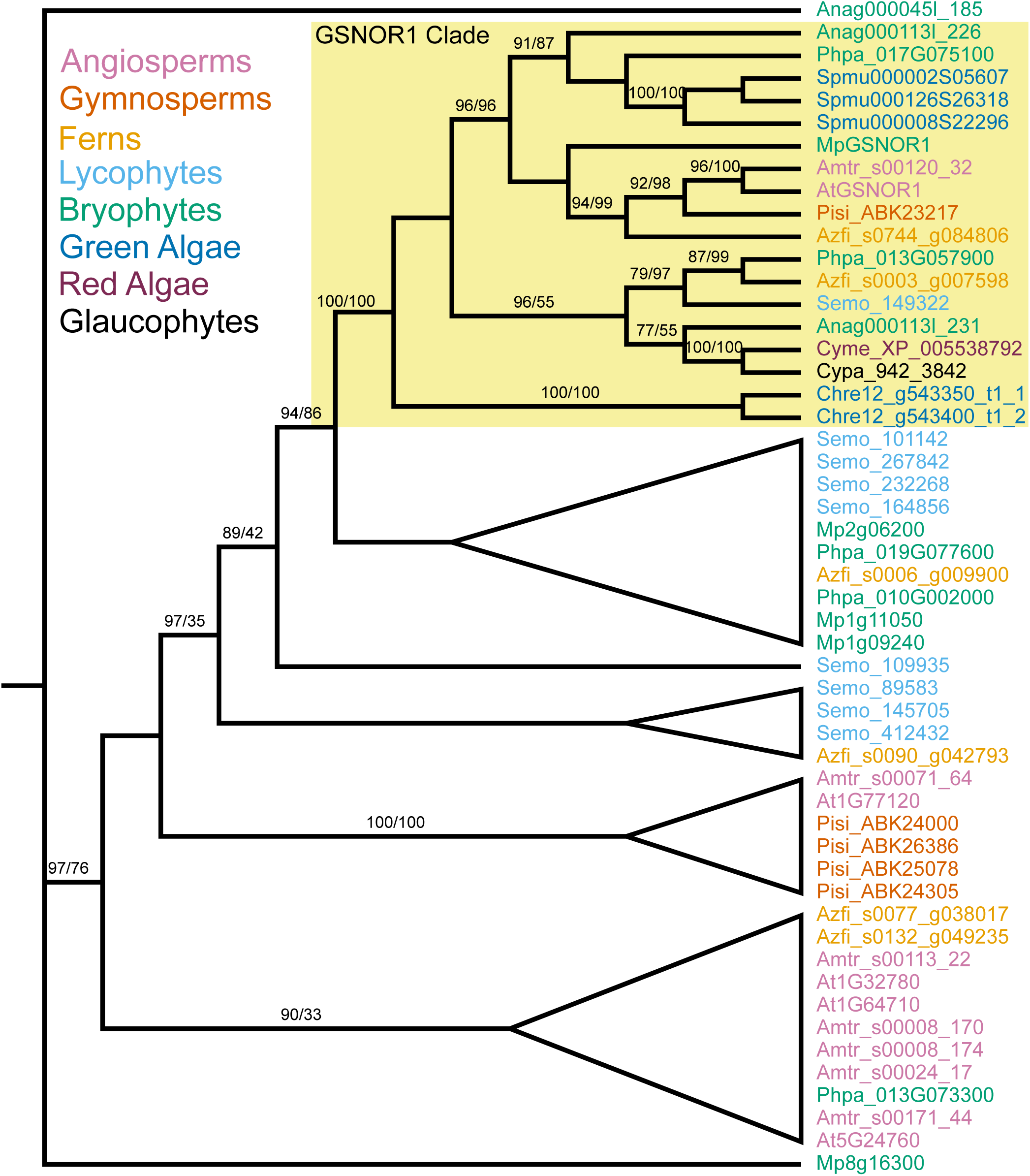
Phylogenetic tree of GSNOR1 and related proteins. Unrooted Maximum likelihood tree created with IQTree and shown as a cladogram. Branches with ultrafast bootstrap and SH-aLRT branch support values exceeding 95% and 80% have high confidence. Ultrafast bootstrap values <75% not shown. The strongly supported GSNOR1 clade is highlighted, other clades shown as cartoons. Species abbreviations: At *Arabidopsis thaliana;* Amtr *Amborella trichopoda*; Pisi *Picea sitchensis;* Azfi *Azolla filiculoides*; Semo *Selaginella moellendorffii*; Anag *Anthoceros agrestis*; Phpa *Physcomitrella patens*; Mp *Marchantia polymorpha*; Spmu *Spirogloea muscicola*; Chre *Chlamydomonas reinhardtii;* Cyme *Cyanidioschyzon merolae*; Cypa *Cyanophora paradoxica*. The different taxonomic groupings are coloured as indicated top left.

Structural analysis using SWISS-MODEL and PyMOL showed that Mp*GSNOR1* also retains the non-zinc-coordinating cysteine residues found in At*GSNOR1 (*Figure 2). This finding further supports its evolutionary stability. The presence of these conserved structural elements, along with its classification in the medium-chain dehydrogenase/reductase (MDR) enzyme family, suggests that Mp*GSNOR1* functions similarly to At*GSNOR1* in maintaining cellular redox balance.

**Figure 2.**
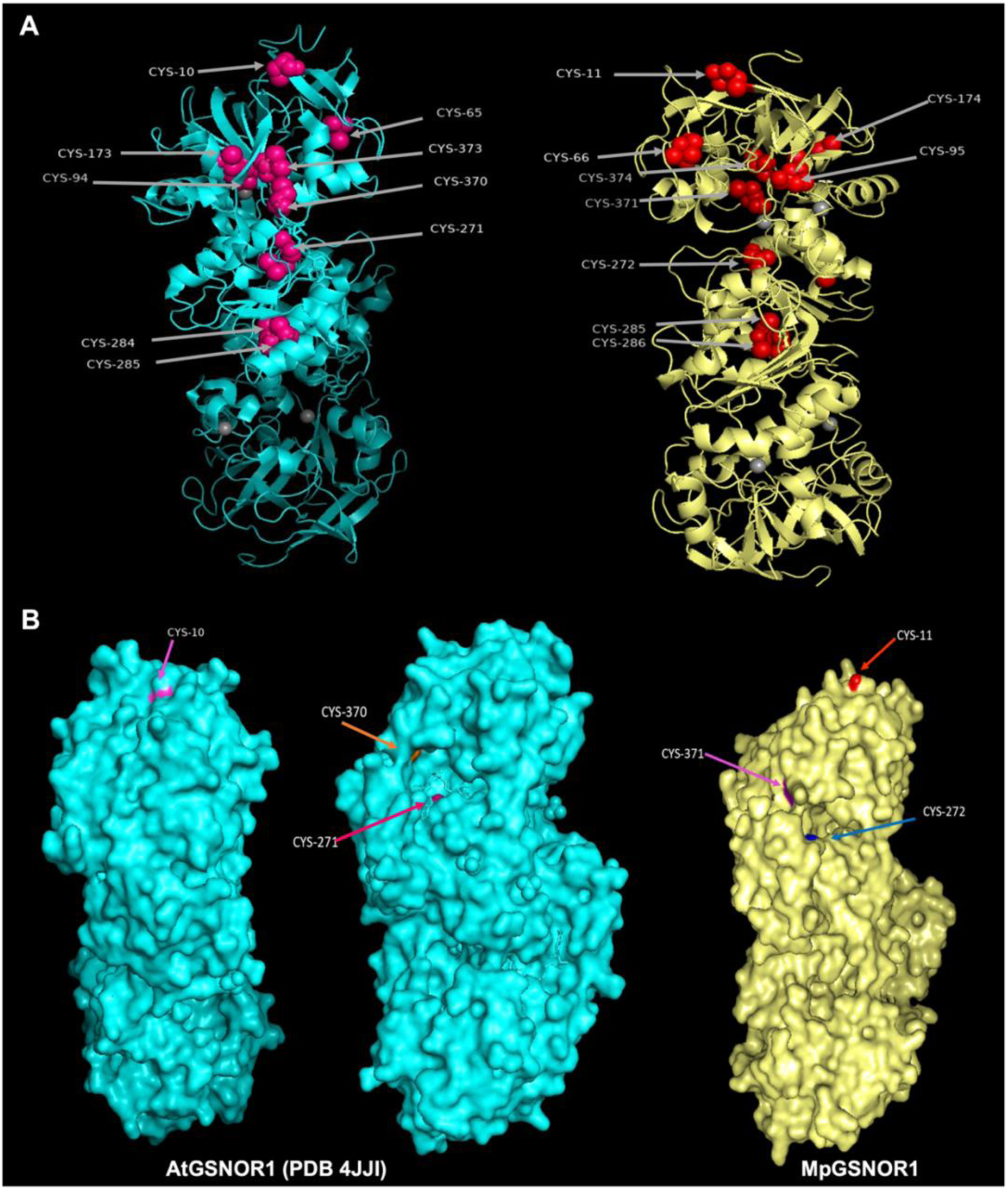
Conservation and solvent accessibility of Cysteine Residues in GSNOR1 from Arabidopsis and Marchantia. (A) Modelled structure of AtGSNOR1 (PDB 4JJI) (blue) and MpGSNOR1 (generated using SWISS-MODEL) (yellow), highlighting conserved active site cysteine residues. (B) Ex-zinc cysteines are annotated, and color coding indicates specific Cys residues: Cys-10 (Pink), Cys-271 (Purple), and Cys-370 (Orange) in AtGSNOR1; Cys-11 (Red), Cys-272 (Blue), and Cys-371 (Violet) in MpGSNOR1. The protein models are oriented to emphasize solvent accessibility, illustrating the relative surface exposure and conservation of these critical residues across species. Images were generated in PyMOL. These visualizations demonstrate the preservation of these residues and highlight their potential importance for GSNOR1 function in diverse species

Overall, these findings suggest that GSNOR1 was present in the last common ancestor of the land plants and has been conserved in Marchantia as a single copy gene. The strong similarity between GSNOR1 in Arabidopsis and Marchantia, along with their shared structural features and key functional residues, suggests potential biochemical and functional similarities.

### Generation of independent Mp*gsnor1* loss-of-function mutants by gene editing

To investigate the functional role of Mp*GSNOR1*, we generated loss-of-function mutants using CRISPR/Cas9-mediated genome editing. Two guide RNA (sgRNA1 and 2) target sites were designed within the second exon of Mp*GSNOR1* (Mp1g16170) and selected for high specificity and minimal off-target effects (Figure S2, Table S1). Transgenic lines were generated through *Agrobacterium-*mediated transformation of wild-type (Tak1) gemmae, and multiple deletion alleles were identified: Mp*gsnor1-1^ge^*(1 bp deletion) and Mp*gsnor1-4^ge^* (4 bp deletion) were independently created using sgRNA 1 and 2 respectively and were selected for further characterisation. Both mutations caused frameshift alterations, leading to premature stop codons that are predicted to inactivate Mp*GSNOR1* (Figure S2).

### Mp*GSNOR1* regulates growth and development in Marchantia

The Mp*gsnor1* loss-of-function mutations significantly impaired growth and development. When cultivated on Gamborg’s B5 medium, both mutants exhibited a stunted phenotype characterised by reduced thallus size and altered growth morphology. Comparing Mp*gsnor1-1^ge^* and Mp*gsnor1-4^ge^* to the wild-type progenitor strain (Tak1), the thallus area decreased by 20.06% (p < 0.05) and 26.98% (p < 0.01), respectively (Figure 3, panel A). After four weeks, the wild-type thallus looked flatter and more dispersed, while the thallus lobes of the Mp*GSNOR1* mutant plants were densely arranged and twisted upward. A significant morphological divergence was also observed in soil-grown plants, with wild-type thallus lobes significantly (p < 0.01) longer than those of the mutants (Figure 3, panel C). Complementation of the Mp*gsnor1-4^ge^* mutant with the Arabidopsis At*GSNOR1* gene (*_pro_35S*:At*GSNOR1*^ge^) in background efficiently restored the thallus growth abnormalities observed in the mutant lines. Thallus area in the complemented lines approached 90% of wild-type levels (Figure 3, panel E). The growth morphology of complemented lines was also nearly identical to that of the wild-type, indicating that the Mp*gsnor1* mutant phenotype was complemented.

**Figure 3.**
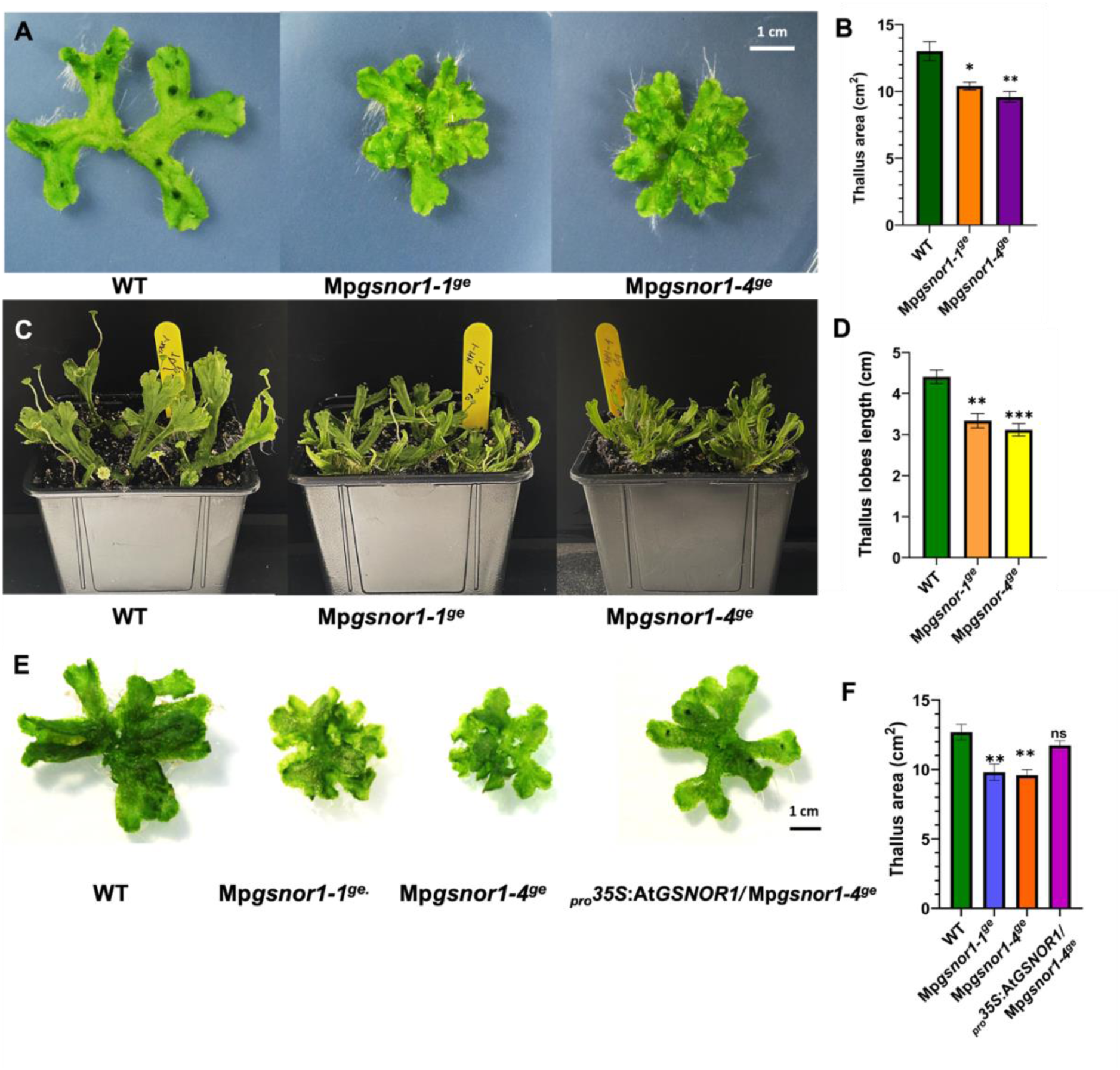
Mp*GSNOR1* loss-of-function and complementation phenotypes in *Marchantia polymorpha.* (A) Representative images of three-to four-week-old wild-type (WT,Tak-1) and Mp*GSNOR1* mutant lines (Mp*gsnor1-1^ge^* and Mp*gsnor1-4^ge^*) grown on ½× B5 Gamborg’s medium from gemmae, showing reduced thallus area in mutants. (C) Eight-week-old plants grown on sterilised soil highlighting reduced leaf size in mutants. Quantification of thallus area (cm²) and thallus lobes length (cm) is shown; error bars represent SEM (n = 5). (B, D) Statistical significance compared to WT is indicated (*p < 0.05, **p < 0.01, unpaired Student’s t-test), with mutants showing 24.32% and 29.32% reductions in leaf length, respectively. (E) Complementation of the Mp*gsnor1-4^ge^*mutant by *_pro_35S*:At*GSNOR1^ge^* restores thallus growth to near WT levels, confirming conservation of GSNOR1 biochemical activity between Marchantia and Arabidopsis. Plants (four weeks old) were grown on ½× B5 Gamborg’s medium from gemmae. (F) Thallus area (cm²) quantification is shown; error bars represent SEM (n = 5). Statistical significance is indicated (ns = non-significant, *p < 0.05, **p < 0.01, unpaired Student’s t-test). Scale bars = 1 cm for all panels.

### Mp*GSNOR1* promotes the growth of rhizoids

Rhizoids are long, tubular protrusions that emerge from epidermal cells on the abaxial surface of the thallus, and primarily function in anchorage. There are two morphologically distinct types of rhizoids, smooth and pegged, with the latter specialised for water transport. (Duckett et al., 2014; Lu et al., 2024). Notably, rhizoid development in bryophytes and root hair formation in angiosperms share a conserved genetic basis, with RSL class I basic helix–loop–helix transcription factors controlling the development of these specialised single-celled structures in the last common ancestor of land plants (Proust et al., 2016). Rhizoid length in Mp*gsnor1* mutants was significantly (p < 0.001) reduced by 46 - 52% compared to wild-type (Figure 4, panel A-E). Like wild-type, mutant plants formed both smooth and pegged rhizoids (data not shown). Importantly, in the complementation line *_pro_35S*:At*GSNOR1/* Mp*gsnor1-4^ge^*, rhizoid development was restored (Figure 4, panel C). Approximately 92% of the rhizoid length lost in the mutants was recovered, indicating that *Arabidopsis GSNOR1* can compensate for Mp*GSNOR1* deficiency. This finding demonstrates the functional equivalence of these genes, indicating that the observed defects in Mp*GSNOR1* mutants are specifically attributable to disruption of Mp*GSNOR1* rather than off-target effects. Furthermore, these results support the high degree of functional conservation of GSNOR1 across species.

**Figure 4.**
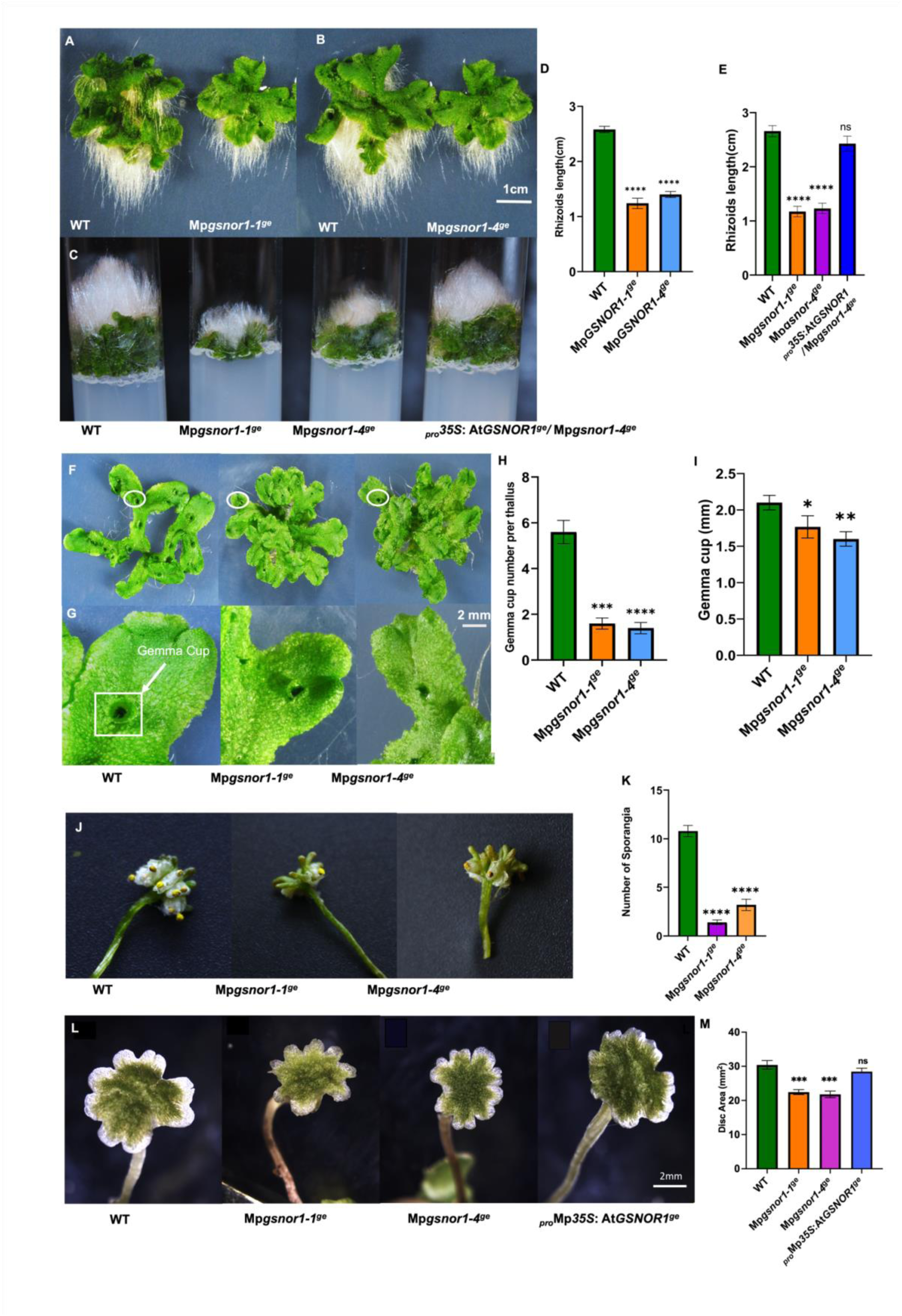
Phenotypic characterisation of Mp*GSNOR1* loss-of-function mutants and complementation lines in *Marchantia polymorpha*. (A–E) Rhizoid development: Representative images of wild-type (WT, Tak-1), Mp*gsnor1-1^ge^* and Mp*gsnor1-4^ge^* mutant lines, and *_pro_35S*:At*GSNOR1^ge^*/Mp*gsnor1-4^ge^* complementation lines (on Mp*gsnor1-4^ge^* background) grown for four weeks on ½× B5 Gamborg’s medium. (A, B) Plants grown on vertical plates showing rhizoid development; (C) plants grown on inverted plates, where negatively phototrophic rhizoids grow away from the light and into the air, highlighting growth differences and recovery upon complementation; (D,E) Quantification of rhizoid length (mean ± SEM, n = 5) in wild-type, mutant, and *_pro_35S*:At*GSNOR1^ge^*/Mp*gsnor1-4^ge^* lines, with significance vs. wild-type (ns, ****p < 0.001) and scale bars = 1 cm (A–C).. (F-I) Gemma cup development: (F,G) Representative images and quantification of gemma cup number per thallus and gemma cup diameter in WT, mutant lines showing reduced gemma formation and size in mutants. (H, I) Bars show average gemma cup number and diameter (mm) ± SEM (n = 5), with significance vs. wild-type (*p < 0.05, **p < 0.01) and scale bars = 2 mm. (J) Sporophyte formation: Representative images and quantification of sporophytes with closed sporangia in WT and mutant lines grown on ½× B5 Gamborg’s medium, showing a significant reduction in sporangia number in mutant lines. (K) Bars show mean ± SEM (n = 5), with significance vs. WT (ns, *p < 0.05, **p < 0.01, ***p < 0.001, ****p < 0.0001) and scale bars = 1 cm (A–C), 2 mm (E–H). (L) Antheridial disc development: Antheridia formation in WT occurs at 3–4 weeks post-germination, whereas mutant lines show delayed initiation (6–7 weeks) and reduced disc size (26.07% and 28.39% smaller, respectively). The complementation line restores both timing and disc size to near WT levels. (M)Bars show mean antheridial disc dimensions ± SEM (n = 5), with significance vs. wild-type (ns, ***p < 0.001) and scale bars = 2 mm.

### Mp*GSNOR1* loss can lead to reduced and delayed development of vegetative reproductive structures

Gemmae, formed within gemma cups on the dorsal thallus, enable clonal propagation in Marchantia. In Mp*GSNOR1* loss-of-function mutants, gemma cup formation was significantly impaired. Compared to wild-type, gemma cup production was reduced by 72% and 76%, respectively, after four to five weeks in growth media (Figure 4, panel F). Mutant plants exhibited delayed gemma cup initiation, reduced gemma cup size, and slower maturation rates. Statistical analysis confirmed these differences are significant (*p* < 0.001). These findings suggest that Mp*GSNOR1* plays a crucial role in regulating asexual reproductive development.

Complementation with *pro35S:*At*GSNOR1^ge^*/Mp*gsnor1-4^ge^*restored gemma formation, reaching 89% of wild-type levels (Figure S3). Gemma cups in the complemented line also recovered to approximately 93% of wild-type size, closely resembling normal development.

### Mp*GSNOR1* promotes the development of sexual reproductive organs

The development of sexual reproductive structures was significantly delayed in Mp*GSNOR1* mutants. Exposure to far-red enriched light (730 nm) induced antheridia formation in wild-type plants within 3–4 weeks of treatment, whereas Mp*gsnor1-1^ge^*and Mp*gsnor1-4^ge^* mutants exhibited a pronounced delay, with initiation occurring only after 6–7 weeks. Additionally, mutant plants developed smaller antheridial discs compared to wild-type. The average diameter of antheridial discs in Mp*gsnor1-1^ge^* and Mp*gsnor1-4^ge^* was distinctly smaller than wild-type (Figure 4, panel L).

Complementation with At*GSNOR1* restored reproductive defects. The complementation line showed a recovery of antheridia formation timing and antheridial disc size, reaching 90% of wild-type levels (Figure 4, panel J). Mp*GSNOR1* therefore promotes the initiation and development of reproductive structures, underscoring its role in controlling the switch from vegetative to reproductive growth.

Crosses between mutant (Mp*gsnor1-1^ge^* and Mp*gsnor1-4^ge^*) and wild-type (*Tak-2*) female plants revealed a reduction in sporophyte formation (Figure 4, panel J) with Mp*gsnor1-1^ge^* and Mp*gsnor1-4^ge^* siring only 87% and 70% of sporophytes compared to wild-type males, respectively (*p* < 0.001). In some cases, sporophyte formation was minimal despite repeated crossings suggesting that male fertility was impaired.

### Mp*gsnor1* mutations reduce GSNOR1 enzymatic Activity

To assess the impact of Mp*GSNOR1* mutations on enzyme activity, we assayed extracts from mutants and compared them to wild-type plants. Both mutant alleles showed a significant reduction in GSNOR1 activity (Figure 5, panel A). To determine if *Arabidopsis* GSNOR1 can compensate for the loss of function in Mp*GSNOR1* mutants, we analysed the enzymatic activity in the *_pro_35S*:At*GSNOR1/* Mp*gsnor1-4^ge^* complementation line. The complementation line exhibited increased GSNOR1 activity relative to wild type confirming that At*GSNOR1* effectively restores enzymatic function in Mp*GSNOR1* mutants (Figure 5A).

**Figure 5:**
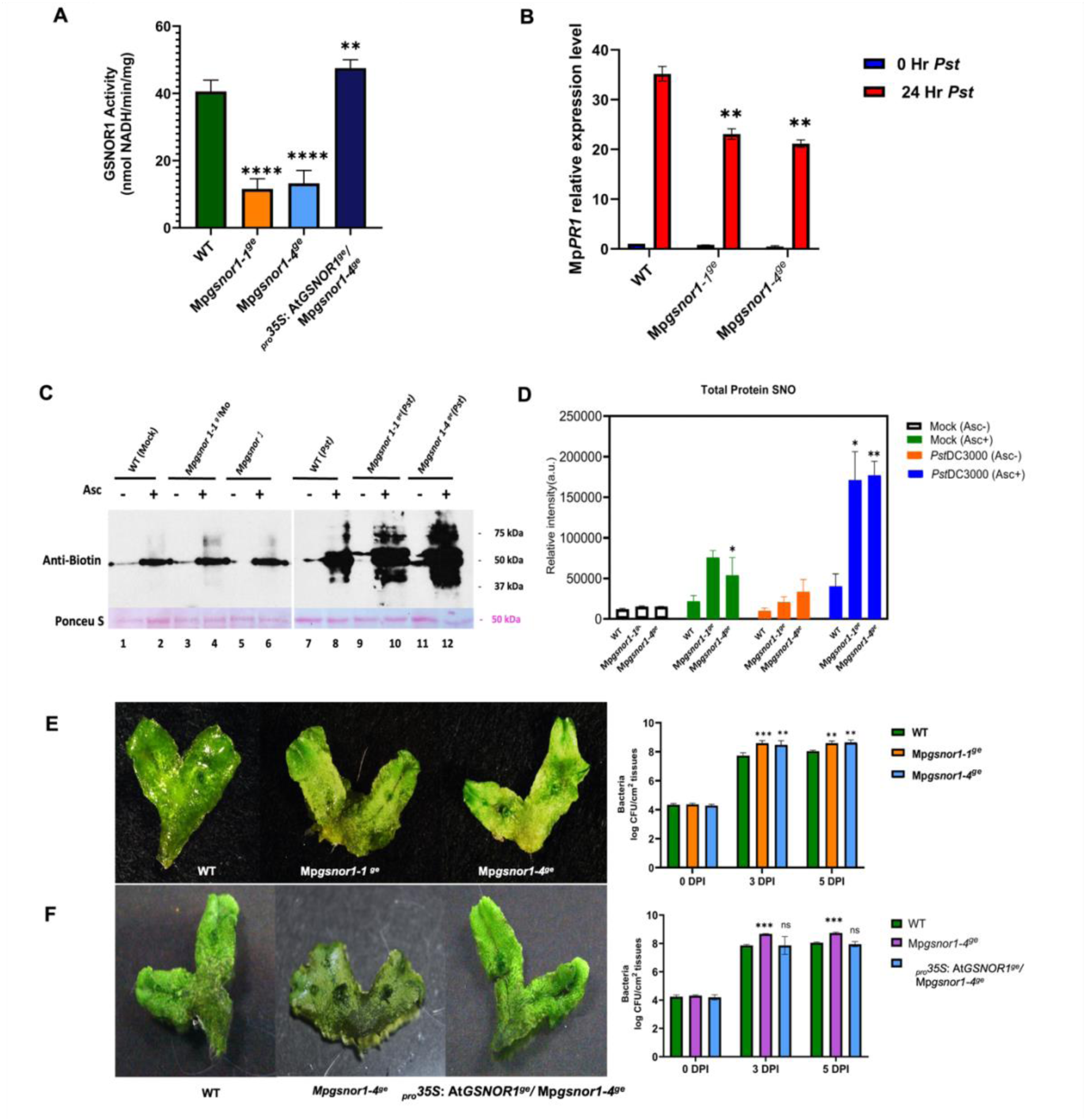
Comparative analysis of GSNOR1 enzymatic activity, total S-nitrosothiol (SNO) levels, and disease resistance in Marchantia wild-type, Mp*gsnor1* mutants, and complementation line. (A) GSNOR1 enzymatic activity measured as NADH oxidation rate per mg protein and expressed relative to wild-type (WT, Tak-1). Mp*gsnor1-1^ge^* and Mp*gsnor1-4^ge^* mutant lines, exhibited reduced activity (29% and 33% of WT), while the *_pro_35S*:At*GSNOR1^ge^*/Mp*gsnor1-4^g^* complementation line restored activity to 120% of WT. Error bars show SEM (n = 5); significance compared to wild-type: **p < 0.01, ****p < 0.001 (unpaired Student’s t-test). (B) *MpPR1* expression in wild-type and *Mpgsnor1* mutants 24 h after *Pst* DC3000 challenged, quantified by qRT-PCR and shown as fold-change relative to mock (0 h). *MpPR1* expression was strongly induced in wild-type plants 24 h post-*Pst* DC3000 infection but markedly reduced in mutants, indicating impaired SA-dependent defence. Asterisks indicate significance relative to the respective 0 h to 24 h treated controls. Error bars = SEM (n = 3); *p < 0.05 compared to 0 h. (C) Total SNO levels, measured by biotin-switch assay in WT and *MpGSNOR1* mutants after *Pst* DC3000 or mock inoculation, were significantly higher in *Pst*-treated mutants than in WT. (D) Quantification of total *S*-nitrosothiol (SNO) levels. Protein extracts from WT and Mp*gsnor1* mutants were analysed following mock or Pst DC3000 inoculation. Ascorbate (ASC) was included as a control in the biotin-switch assay. Signal intensities were quantified with ImageJ. Bar graph: white, mock/–ASC; green, mock/+ASC; orange, Pst/–ASC; blue, Pst/+ASC. Error bars = SEM (n = 3), *p < 0.05, **p < 0.01 (unpaired Student’s t-test). (E, F) Disease symptoms and bacterial growth (CFU cm⁻²) in WT, mutant lines and complementation line at 0, 3, and 5 days post-inoculation with *Pst* DC3000. Mutants showed enhanced susceptibility, while complementation restored resistance. Error bars 0 h controls: ns = non-significant *p < 0.05, ***p < 0.001,; unpaired Student’s t-test.

These findings highlight the essential role of Mp*GSNOR1* in regulating nitrosative stress and demonstrate functional conservation between *Marchantia* and *Arabidopsis* GSNOR1 in maintaining redox homeostasis.

### Role of Mp*GSNOR1* in SNO homeostasis and immunity

To determine the role of MpGSNOR1 in immunity, we first characterised SNO levels in mutant and wild-type plants following infection by *Pst* DC3000. Plants were treated with a bacterial suspension of virulent *Pst* DC3000 or mock treated with a 10 mM MgCl₂ solution as a control, then thallus tissue samples were collected 24 hours post-inoculation, and total SNO levels were determined through the biotin switch assay (Jaffrey & Snyder 2001). The results were striking: *Pst* DC3000-challenged plants, exhibited a significant increase in SNO levels compared to mock treated plants, but the increase was much greater in the mutants than in wild type (Fig. 5C, D). This suggests that the absence of Mp*GSNOR1* compromises the regulation of *S*-nitrosylation, leading to a hyper-accumulation of SNOs. The increase in SNO activity highlights the role of *Pst* DC3000 in inducing an initial surge of NO, which subsequently facilitates elevated SNO production. This process may contribute to immune signalling in Marchantia, further supporting the regulatory role of Mp*GSNOR1* in plant immunity

We next investigated the role of Mp*GSNOR1* in basal disease resistance, by assessing the susceptibility of Mp*gsnor1-1^ge^* and Mp*gsnor1-4^ge^* mutants to *Pseudomonas syringae* pv. tomato (*Pst*) DC3000. Three-week-old thalli from wild-type and mutant plants were inoculated with *Pst* DC3000 at a concentration of 10□ cfu/mL. Bacterial proliferation was quantified at zero, three and five days post-inoculation following the protocol described by Gimenez-Ibanez et al., 2019. Both mutants exhibited significantly increased bacterial growth compared to wild-type plants (Figure 5E). At three and five days post-inoculation, bacterial densities in the mutants reached 10^8^–10^9^ cfu/cm², whereas wild-type plants restricted growth to approximately 10^7^ cfu/cm². Symptoms were most pronounced at the basal regions of the thalli, although bacterial colonisation was also observed at the apical notches. A progressive increase in susceptibility was evident over time, with more severe symptoms by day five.To determine whether Arabidopsis *GSNOR1* could restore *Pst* DC3000 resistance, the *_pro_35S*:At*GSNOR1*^ge^/ Mp*gsnor1-4^ge^* complementation line was analysed. This line showed enhanced resistance, correlating with reduced bacterial loads and attenuated disease symptoms (Figure 5F). These results suggest that Mp*GSNOR1* is essential for basal resistance to *Pst* Dc3000. The functional complementation by At*GSNOR1* confirms its conserved role in immune responses. Our findings suggest that GSNOR1-mediated redox homeostasis is a pivotal factor in plant defense mechanisms across species.

### Mp*gsnor1* compromises SA-Regulated gene expression

SA is a key defence hormone, essential for plant immunity against *Pst* DC3000. While primarily studied in angiosperms, SA signalling is evolutionarily conserved in bryophytes, including Marchanti*a* (Gimenez-Ianez et al., 2019). To investigate the role of Mp*GSNOR1* in SA-mediated defence, we analysed the expression of SA-related genes in wild-type and Mp*GSNOR1* mutant plants following *Pst* DC3000 infection. Quantitative Real Time RT-PCR analysis (Figure 5B) revealed significantly reduced expression of *Pathogenesis-Related 1* (*PR1*), a marker of SA signalling, in Mp*gsnor1-1^ge^*and Mp*gsnor1-4^ge^* mutants compared to wild-type plants 24 hours post-infection indicating impaired SA-dependent defence activation.

These findings suggest that Mp*GSNOR1* regulates SA-related gene expression, playing a critical role in plant immunity. The reduced induction of *PR1* in Mp*GSNOR1* mutants highlights its importance in SA biosynthesis and signalling, reinforcing its function in basal disease resistance against *Pst* DC3000.

## DISCUSSION

### GSNOR1 as an evolutionarily conserved regulator of NO signaling

Our study showed that GSNOR1 has a conserved and indispensable role in NO homeostasis, development, and immunity in Marchantia. Our phylogenetic analysis indicates that GSNOR is conserved in all lineages of the Archaeplastida, illustrating the importance of this enzyme for core biological processes. GSNOR was present in the common ancestor of land plants and retained in Marchantia, where it is encoded by a single gene Mp*GSNOR1*, the orthologue of Arabidopsis At*GSNOR1*. The two proteins share extensive sequence and structural similarity throughout their lengths, highlighting the evolutionary constraint on this enzyme and its non-redundant function in plant biology (Lynch & Conery, 2000; Xu et al., 2013). In addition to this, the structural and functional similarities help us better understand the plant immune system and its evolutionary development.

### Functional impact and compensatory pathways in Mp*GSNOR1* mutants

We found that although GSNOR enzyme activity is strongly reduced in Mp*gsnor1-1^ge^* and Mp*gsnor1-4^ge^* mutants to 29–33% of wild-type levels, it is not completely eliminated. This residual activity is similar to what was observed in Arabidopsis At*GSNOR1* mutants (Feechan et al., 2005) and suggests that alternative pathways can partially compensate for the loss of GSNOR activity. Recent biochemical studies highlight that aldo-keto reductases (AKRs) may play a role in GSNO metabolism during stress (Stomberski et al., 2019; Treffon & Vierling, 2022), although there is not genetic evidence to support functional compensation of GSNOR by aldo-keto reductases in planta. In Arabidopsis, AKR4C8 and similar members of its clade show enzymatic properties comparable to the functions of mammalian AKR1A1 and can reduce GSNO albeit with less catalytic efficiency and specificity than GSNOR (Treffon et al., 2021; Treffon & Vierling, 2022). Therefore, one possibility is that AKRs or other enzymes work in part to replace GSNOR activity in the control of GSNO and nitrosative stress in MpG*SNOR1* mutants. These alternative pathways need more biochemical and genetic investigation.

### Nitric oxide homeostasis and the many functions of GSNOR1 in Marchantia development

NO is a central signaling molecule in plants, with *S*-nitrosylation modulating key physiological processes and stress responses (Allain et al., 2011; Besson-Bard, Courtois, et al., 2008; Besson-Bard, Pugin, et al., 2008; Gaupels et al., 2011; Melotto et al., 2006). In our examination of CRISPR/Cas9-induced Mp*GSNOR1* mutants, we found that there is a significant alteration of the vegetative and reproductive development. These included smaller plant size, smaller thalli, and different thallus shapes. The mutants had shorter and fewer rhizoids, along with significant delays and reductions in sexual structures such as antheridia and sporophytes. These alterations were similar to those observed in the developmental and reproductive structures in Arabidopsis *GSNOR1* mutants (Kwon et al., 2012). These findings point to the critical role of GSNOR1 in reproductive development, likely through NO-regulated pathways that control organ formation and light-responsive transitions (Naramoto et al., 2019).

These phenotypic abnormalities may reflect perturbations in hormonal pathways. Mp*GSNOR1* mutations likely impact auxin, ethylene, and cytokinin pathways that are essential for Marchantia development (Aki, Mikami, et al., 2019; Katayose et al., 2021; Kato et al., 2017; Pande et al., 2022). For instance, NO-mediated *S*-nitrosylation has been shown to alters the activity of auxin receptors (TIR1/AFBs) and response factors in Arabidopsis (Fernández-Marcos et al., 2011; Iglesias et al., 2018; Pande et al., 2022b; Terrile et al., 2012), and these auxin signalling pathway components are conserved in Marchantia (Eklund et al., 2015; Flores-Sandoval et al., 2015; Kato et al., 2017; Suzuki et al., 2023). Changes in NO balance in Mp*GSNOR1* mutants may therefore disrupt auxin signalling, which affects growth. Similarly, *S*-nitrosylation impacts the biosynthesis and action of ethylene. Enzymes like ACC synthase and ACC oxidase, along with their regulators, are sensitive to NO status (Li et al., 2022; Pattyn et al., 2021). These pathways might be further explored using ethylene-specific inhibitors of biosynthesis and perception (Depaepe & Van Der Straeten, 2016; Li et al., 2022; Lindermayr et al., 2006; Qi et al., 2022)

Cytokinin signaling is crucial for gemma cup formation and is regulated by genes like Mp*RRB* and Mp*GCAM1* (Aki et al., 2019, 2022). Imbalances between auxin and cytokinin could explain some developmental defects in Mp*gsnor1* mutants. Experimentally, NO has been found to change cytokinin signalling in a directed manner (Feng et al., 2013; Pande et al., 2022). In particular, *S*-nitrosylation of key signalling molecules in Arabidopsis, including HISTIDINE PHOSPHOTRANSFER PROTEIN 1 (AHP1), cleaves the phosphorelay cascade that is central to cytokinin signaling (Feng et al., 2013; Pande et al., 2022). Cytokinin signalling components including AHP1 are conserved in Marchantia, so it is possible that similar regulation by SNO contributes to the effects seen on gemmae cup formation and development.

The traits seen in Mp*GSNOR1* mutants may indicate broader changes in hormonal signalling networks and transcriptional regulation. These changes could affect morphogenesis, rhizoid development, and reproduction. While more hormonal and signalling studies are necessary, these findings suggest that GSNOR1 has a complex role in Marchantia. They also help us understand the shared mechanisms that drive plant growth, development, and adaptation.

### Evolutionary Conservation of Mp*GSNOR1* in Immune Regulation

GSNOR1 plays a key role in maintaining the balance of specific cellular molecules by controlling global *S*-nitrosylation (Feechan et al., 2005; Kwon et al., 2012). In Arabidopsis, the loss of At*GSNOR1* results in impaired GSNO breakdown, leading to higher levels of *S*-nitrosylated proteins and GSNO (Feechan et al., 2005; Tada et al., 2008). *GSNOR1* itself can be modified by *S*-nitrosylation, particularly at the Cys-10 residue, and this change has been shown to reduce its enzymatic activity in both plants and other organisms (Kubienová et al., 2013; Lee et al., 2008; Zhan et al., 2018). These redox modifications of GSNOR1 highlight the complex nature of NO signalling in plant cells.

Studies using reverse genetics have shown that GSNOR1 plays a central role in controlling *S*-nitrosylation and the expression of SA-dependent genes, which in turn shape the immune responses of vascular plants (Feechan et al., 2005; Xia et al., 2014). However, the role of GSNOR1 in non-vascular plants remained largely unknown. By investigating Marchantia (Bowman et al., 2017, 2022; Alcaraz et al., 2018), our study addresses this gap and provides new insights into the evolutionary origins of NO-based immunity.

Our findings demonstrate that Mp*GSNOR1* mutants accumulate higher levels of SNO proteins, indicating that the absence of *GSNOR1* disrupts NO-dependent signalling networks. This supports the findings in angiosperms and supports the hypothesis that SNOs act as mediators of NO bioactivity in response to pathogen challenge, and SNOs playing complementary roles in plant disease resistance (Feechan et al., 2005; Hussain et al., 2019a; Lee et al., 2008; Tada et al., 2008).

Phytohormones such as SA and jasmonate are central to immune regulation in both Marchantia and angiosperms (Boatwright & Pajerowska-Mukhtar, 2013; Dempsey et al., 2011; Gimenez-Ibanez et al., 2019; Monte, 2023). Exposure to pathogens results in upregulation of SA marker genes in Marchantia, indicative of an ancient role for SA-mediated immunity(Gimenez-Ibanez et al., 2019). Consistently, Mp*GSNOR1* mutants show reduced SA-dependent gene expression, including lower Mp*PR1* levels, highlighting a conserved role for GSNOR1 in modulating SA signalling (Feechan et al., 2005). Interestingly, the perception of SA in Marchantia appears to differ from that in Arabidopsis, at least as regards immunity. In Arabidopsis, SA signalling is mediated by the receptor NPR1 together with its cofactor TGA1 (Després et al., 2003; Shearer et al., 2012), so that SA induced transcriptional reprogramming is lost in the *npr1* mutants. Marchantia contains single copies of *NPR*1 and *TGA1*, suggesting that NPR-dependent SA signalling was present in the most recent common ancestor of land plants (Jia et al., 2023). Functional studies show that although MpNPR1 can bind SA, mutants are hypersensitive to SA and the transcriptional reprogramming that occurs in response to SA is largely intact in Mp*npr1* mutants, implying the existence of an alternative SA sensing pathway in Marchantia (Jeon et al, 2024). In addition, although S-nitrosylation of *NPR1* at Cys-156 by GSNO promotes its oligomerisation and stabilises protein homeostasis in response to SA in Arabidopsis (Tada et al., 2008) comparative sequence analysis of 194 NPR proteins found that Cys156 is conserved only in Brassicaceae so it is not clear whether MpNPR1 is regulated by *S*-nitrosylation of cysteine residues (Jeon et al., 2024). It will be interesting to determine what the alternative SA signalling pathways are and whether their components are SNO regulated.

The bacterial pathogen *Pst* DC3000 has been identified as a virulent agent in Marchantia (Gimenez-Ibanez et al., 2019). Our results show that Mp*GSNOR1* mutants are more susceptible to *Pst* DC3000, whereas complementation with At*GSNOR1* restores resistance. This functional conservation underscores the broad importance of *GSNOR1* across land plants, as Arabidopsis *GSNOR1* mutants similarly exhibit increased pathogen susceptibility (Feechan et al., 2005). The cross-species complementation of Mp*GSNOR1* mutants with At*GSNOR1*, resulting in restored pathogen resistance, provides direct evidence for the evolutionary conservation of GSNOR1-mediated immune regulation.

## Conclusions and Future Directions

Our study establishes GSNOR1 as a central, evolutionarily conserved regulator of NO homeostasis, hormone signaling, development, and immunity in Marchantia (Figure 6). While the fundamental roles of GSNOR1 are shared across land plants, species-specific adaptations in hormonal and immune responses persist. Future work should aim to biochemically and genetically dissect the compensatory roles of AKRs and elucidate the molecular interplay between GSNOR1 and NPR1 or alternative SA receptors in non-vascular plants, paving the way for targeted improvements in crop resilience and disease resistance.

**Figure 6:**
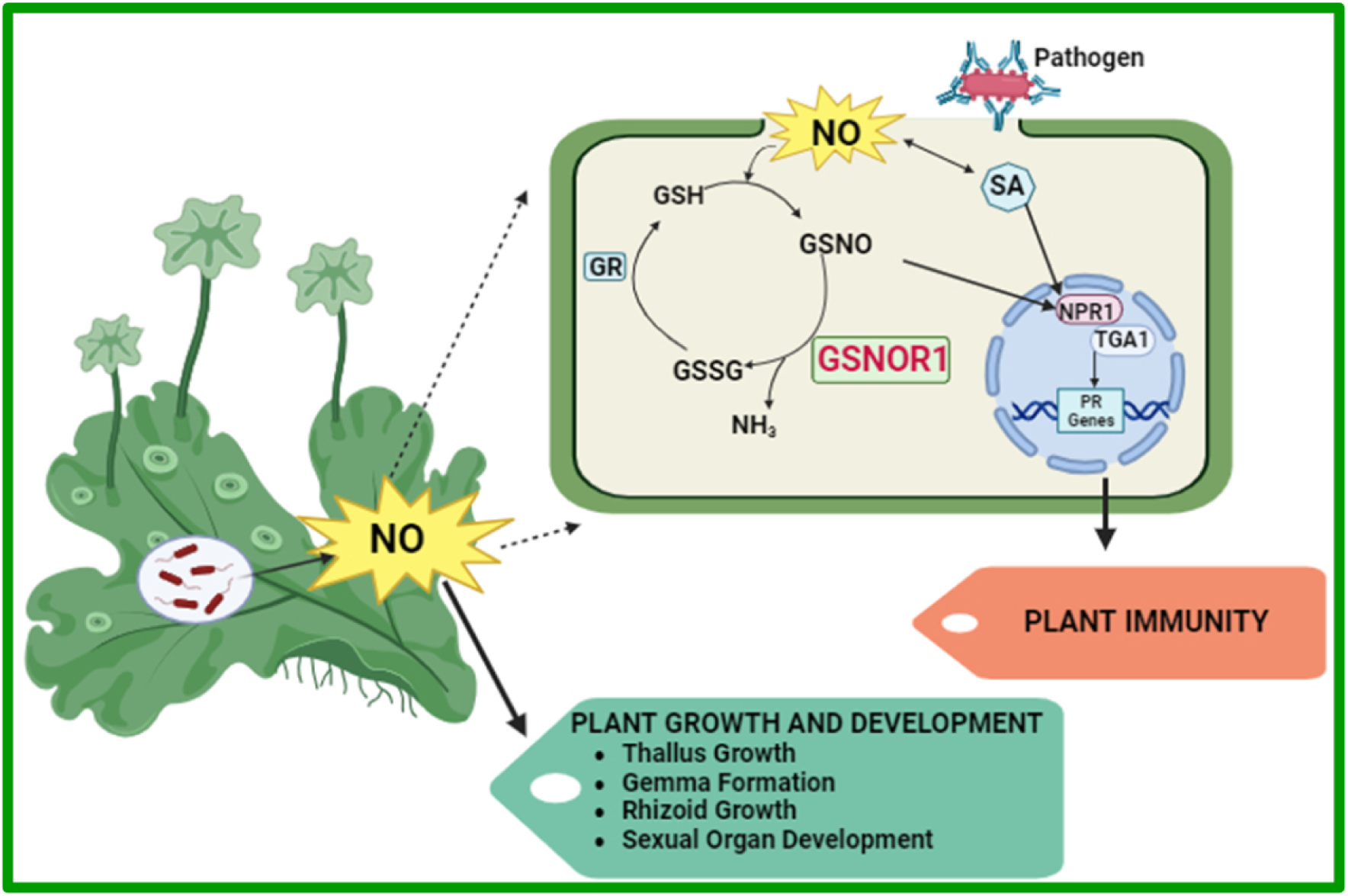
Schematic overview of GSNOR1 function in Marchantia. This diagram outlines the role of GSNOR1 in plant immunity and development in the non-vascular plant Marchantia. It shows how GSNOR1 indirectly regulates SA-mediated defence by controlling NO levels through GSNO degradation. Upon pathogen detection, SA synthesis is triggered, and GSNOR1 modulates this signalling to activate defence genes. The figure also highlights the crosstalk between SA and NO pathways mediated by GSNOR1. Beyond immunity, GSNOR1 influences developmental processes, including thallus growth, gemma formation, rhizoid development, and reproductive organ formation by maintaining NO balance, which is vital for environmental adaptation. Figure generated in BioRender.com.

## Acknowledgments

NT gratefully acknowledge the late GL for his invaluable guidance and inspiration throughout this doctoral work. The project was originally conceived by GL, whose enthusiasm left a lasting impact, and his passing is remembered with deep respect. We also thank the University of Edinburgh, Institute of Molecular Plant Sciences, the Edinburgh Plant Growth Facility, the Wellcome Centre for Cell Biology, and the Centre for Optical Instrumentation Laboratory (COIL) for providing research facilities. NT gives thanks to the University of Dhaka for providing her study leave during this doctoral study.

## Author Contributions

GJ planned the study, GJ, JG and NT designed the experiments. NT performed the experiments under the supervision of GL and JG. JG constructed the phylogenetic tree Image. NT prepared the figures and drafted the manuscript. GL reviewed the initial draft, and JG provided subsequent revisions and edits as required. JG and NT approved the manuscripts.

## Data Availability

All data generated or analysed in this study are available within the published article and its supplementary materials.

## Supplementary Figures

**Figure S1: Conservation of Catalytic, Structural, and Binding Residues in Plant GSNORs.** Multiple sequence alignment of GSNOR proteins from Marchantia, moss, and green algae. Amino acid sequences were aligned using **Clustal Omega** (Sievers et al., 2011) and visualised with **ESPript 3.0** (Robert & Gouet, 2014). Species abbreviations: At – *Arabidopsis thaliana*; Azfi – *Azolla filiculoides*; Phpa – *Physcomitrella patens*; Mp – *Marchantia polymorpha*; Chre – *Chlamydomonas reinhardtii*. Residue functional annotations are based on structural and biochemical studies of *Arabidopsis thaliana* GSNOR (e.g., Kubienová et al., 2013; Leterrier et al., 2011). Black filled circles (●) indicate residues coordinating the catalytic zinc atom; black filled triangles (▴) mark residues coordinating the structural zinc atom; blue filled circles (D) denote residues interacting with the NAD(H) cofactor; green squares (D) indicate substrate-binding residues (GSNO/HMGSH). Asterisks (*) mark solvent-accessible cysteine residues potentially targeted for redox-based post-translational modifications. Orthologues from different species are enclosed within dotted boxes to highlight sequence conservation

**Figure S2. Sanger sequencing chromatograms showing CRISPR/Cas9-generated mutations in Exon 2 of the Mp*GSNOR1* gene.** (A) Schematic representation of the Mp*GSNOR1* gene. White boxes represent untranslated regions (UTRs), while colored boxes indicate protein-coding exons (CDS). Black connecting lines correspond to introns. The positions of the two single-guide RNAs (sgRNA1 and sgRNA2) used for genome editing are shown. (B) Chromatograms displaying the Exon 2 sequence for the wild type (WT), a single-nucleotide deletion mutant (Mp*gsnor1-1^ge^*). These mutations cause frameshifts predicted to disrupt MpGSNOR1 protein function.

**Figure S3: Comparative assessment of gemma cup development and gemma size in *Marchantia* wild-type, mutant, and complementation lines.** The figure compares gemma production and size in the (A) Wild-type, (B) Mp*gsnor1-1^ge^* and (C) Mp*gsnor1-4^ge^* mutants, and (D) the complementation line *_pro_35S*:At*GSNOR1^ge^*/Mp*gsnor1-4*, Differences in gemma formation frequency and size are observed among the lines, with the complementation line showing marked restoration of both traits to near wild-type values. Bars depict mean values ± SEM (n = 5), with statistical significance relative to wild-type indicated (ns = non-significant, *p < 0.05, **p < 0.01, ****p < 0.001; unpaired Student’s t-test). Scale bars = 1 cm.

## Abbreviations

(At*GSNOR1*): Arabidopsis *S*-nitrosoglutathione reductase 1
(EvoMPMI): Evolutionary molecular plant-microbe interactions
(Marchantia, Mp): *Marchantia polymorpha*
(*Pst*): *Pseudomonas syringae*
(SA): salicylic acid
(Cys): cysteine
(SNO): *S*-nitrosothiol
(NO): nitric oxide
(qRT-PCR): Quantitative real-time PCR
(GSNO): *S*-nitrosoglutathione
(GSNOR): *S*-nitrosoglutathione reductase
(Mp*GSNOR*): Marchantia *S*-nitrosoglutathione reductase
(Tak-1): Takaragaike-1
(WT): wild-type

## Supplementary Files

**Table S1-.**
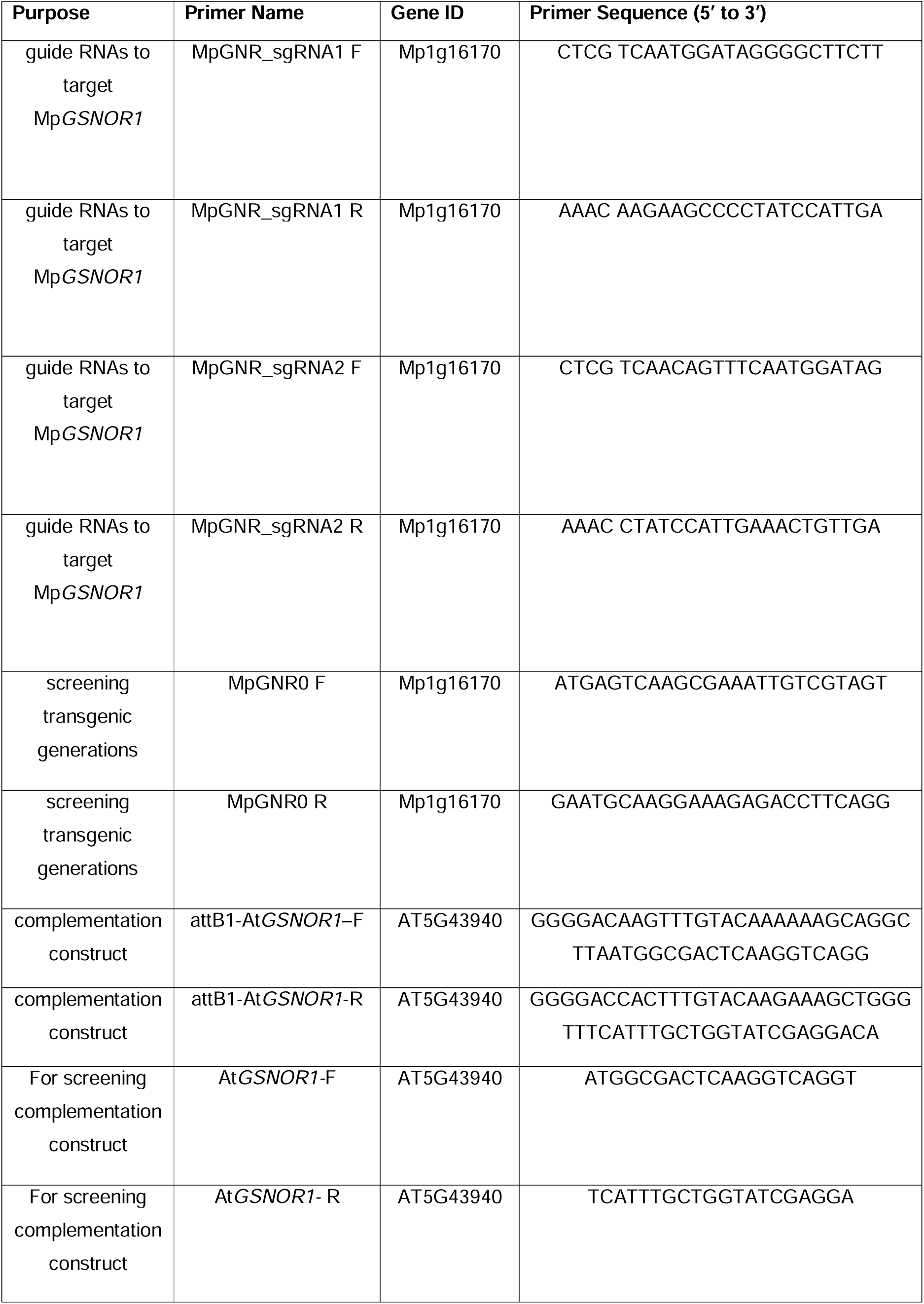
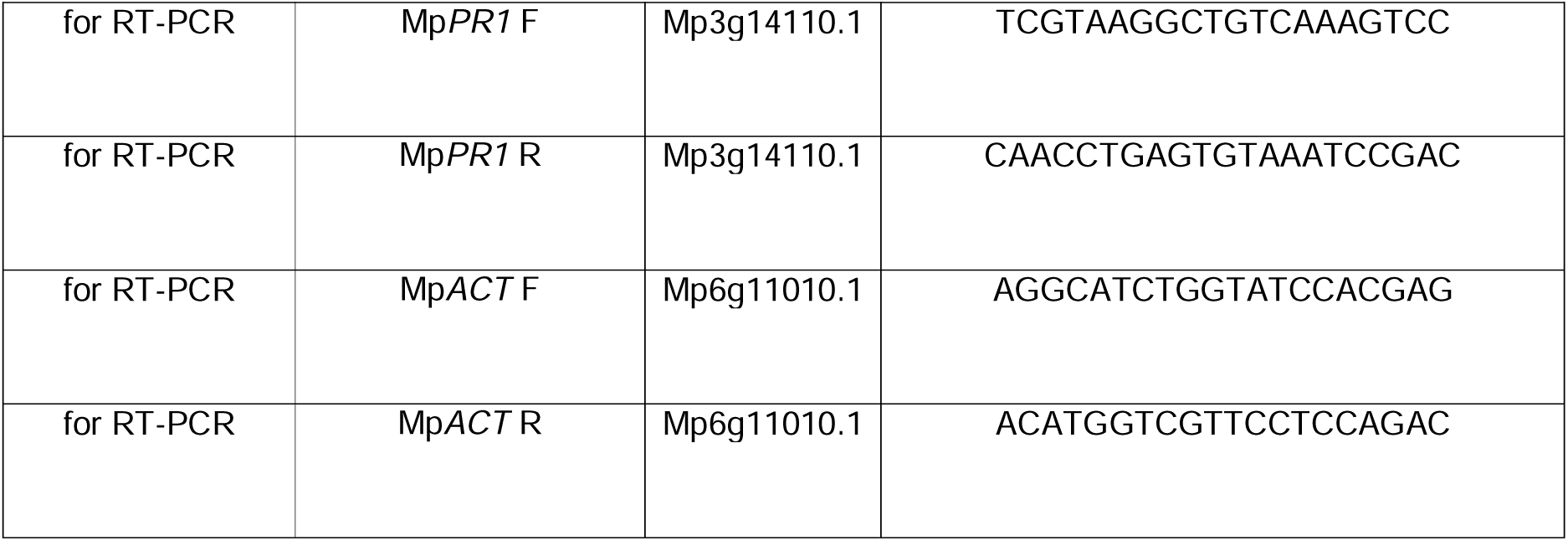
List of primers used for this study.

